# A topographical atlas of αSyn dosage and cell-type expression in the mouse brain and periphery

**DOI:** 10.1101/2023.10.05.559770

**Authors:** Haley M. Geertsma, Zoe A. Fisk, Lillian Sauline, Alice Prigent, Kevin Kurgat, Steve M. Callaghan, aSCENT-PD consortium, Michael X. Henderson, Maxime W.C. Rousseaux

## Abstract

Parkinson’s disease (PD) is the second most common neurodegenerative disease worldwide and presents pathologically with Lewy pathology and dopaminergic neuron loss. Lewy pathology contains aggregated αSynuclein (αSyn), a protein encoded by the *SNCA* gene which is also mutated or duplicated in a subset of familial PD cases. Due to its predominant presynaptic localization, immunostaining for the protein results in diffuse signal, providing little insight into the types of cells expressing αSyn. As a result, insight into αSyn expression-driven cellular vulnerability has been difficult to ascertain. Using a combination of knock-in mice that target αSyn to the nucleus of cells (*Snca^NLS^*) and *in situ* hybridization of *Snca* in wild-type mice, we systematically map the topography and cell types expressing αSyn in the mouse brain, spinal cord, retina, and gut. We find a high degree of correlation between αSyn protein and RNA levels across multiple brain regions and further identify cell types with low and high αSyn. We found that αSyn is highly expressed in neurons, particularly those involved in PD and to a lower extent in non-neuronal cell types, notably those of oligodendrocyte lineage. We also find that αSyn is devoid in certain neuron types (e.g. ChAT-positive motor neurons), and that all enteric neurons express αSyn to a certain degree. Taken together, this atlas provides much-needed insight into the cellular topography of αSyn, and provides a quantitative map to test assumptions about the role of αSyn in network vulnerability in PD and other αSynucleinopathies.

## Introduction

Parkinson’s disease (PD) and related dementias exist along a clinical and pathological continuum, sharing a common protein pathology. αSynuclein (αSyn) undergoes aggregation and is thought to contribute to disease pathogenesis in several neurodegenerative diseases (i.e. PD, Lewy body dementia [LBD], multiple system atrophy [MSA]), collectively termed αSynucleinopathies^1,2^. αSyn accumulates in proteinaceous structures called Lewy bodies and Lewy neurites (collectively: Lewy pathology) except for MSA where it accumulates in glial cytoplasmic inclusions [GCIs])^3,4^. Mutations and copy number gains in the gene encoding αSyn, *SNCA*, can also cause PD^5,6^. Thus, the convergence of pathology and genetics place αSyn at the center of pathogenesis for these disorders. Despite its linkage to PD over 25 years ago, less is known about the native role of αSyn and how it transitions from a native, functional form to a pathogenic state^7^. Normally, αSyn is found at the presynaptic membrane, and to lesser extents in the nucleus, mitochondria, and cytosol^8–12^. In disease conditions, αSyn mislocalizes from the membrane to non-synaptic compartments (e.g. soma and processes) where it adopts unstructured and aggregate-prone conformations, eventually forming intracellular aggregates^13–15^. How this process is linked to the ultimate formation of Lewy pathology remains unclear.

Increasing evidence suggests that expression levels of αSyn, as well as posttranslational and conformational changes, all play a role in modulating its aggregation^5,6,16^. Biochemical and biophysical data have begun to establish key modifications and conformations that lead to αSyn accumulation and aggregation^17–19^. Moreover, increasing data suggest that misfolded αSyn may act in a prionoid manner, spreading from one cell to another, initiating aggregate formation^20,21^. This process is generally thought to be dependent upon the local concentration of native αSyn within the target cell^22–24^. Yet local, topographical dosage of αSyn has been relatively understudied to date. This may be in part due to the limited tools available to glean meaningful insight at sufficient resolution. Since αSyn is mostly a presynaptic protein, immunodetection in healthy individuals or model organisms results in diffuse presynaptic staining, making it difficult to attribute expression to any particular cell. RNA detection methods such as fluorescence *in situ* hybridization have offered insight into this distribution, but there is no quantitative *Snca* map using this technique. Furthermore, it is important to recognize that cellular RNA and protein levels may not always correlate, thereby emphasizing the necessity for a comprehensive brain-wide comparative analysis encompassing both proteomic and transcriptomic dimensions^25^.

We recently developed a *Snca^NLS^* mouse allele to study the consequence of αSyn nuclear mislocalization^26^. These mice have a nuclear localization sequence (NLS) and Flag epitope tag knocked into the C-terminus of the endogenous *Snca* gene that codes for αSyn. While characterizing these mice, we serendipitously discovered that targeting the protein to the nucleus provides unprecedented detail on the cellular and anatomical topography of this predominantly synaptic protein. We took advantage of this cell body localization to map the regional and cellular localization of αSyn throughout the brain on the protein level and compared this to *Snca* RNA. We present a systematic study of its topography throughout the brain, spinal cord, retina, and enteric nervous system. We find that αSyn is broadly expressed in neurons of both the central and enteric nervous system (CNS and ENS, respectively) with specific region and neuron type enrichment. In addition, we find that αSyn is expressed in a subset of Olig2-positive cells, likely representing oligodendrocyte precursor cells. To our surprise, some cell clusters including ChAT-positive cells of the lateral reticular nucleus (LRN) and spinal cord motor neurons do not express αSyn.

## Results

### Whole-brain mapping reveals distinct domains of anatomical αSyn expression and distribution

To study the global topography of αSyn across the mouse brain, we used four parallel approaches: 1) *Snca* RNA mapping in wild-type mice, 2) regional protein mapping with immunofluorescence and immunohistochemical staining of *Snca^NLS^* knockin mice, 3) whole brain clearing and immunolabeling, and 4) cell type marker co-localization.

To determine the cellular expression of αSyn, we first stained tissue for *Snca* mRNA in wildtype mouse brain sections (**Fig 1A**). These sections were then put through a cell classifier in QuPath to classify cells as either high (greater than one standard deviation above the mean), medium (one standard deviation above or below the mean), or low (one standard deviation or more below the mean) *Snca* expression. (Supp **Fig 1A**).

**Figure 1:**
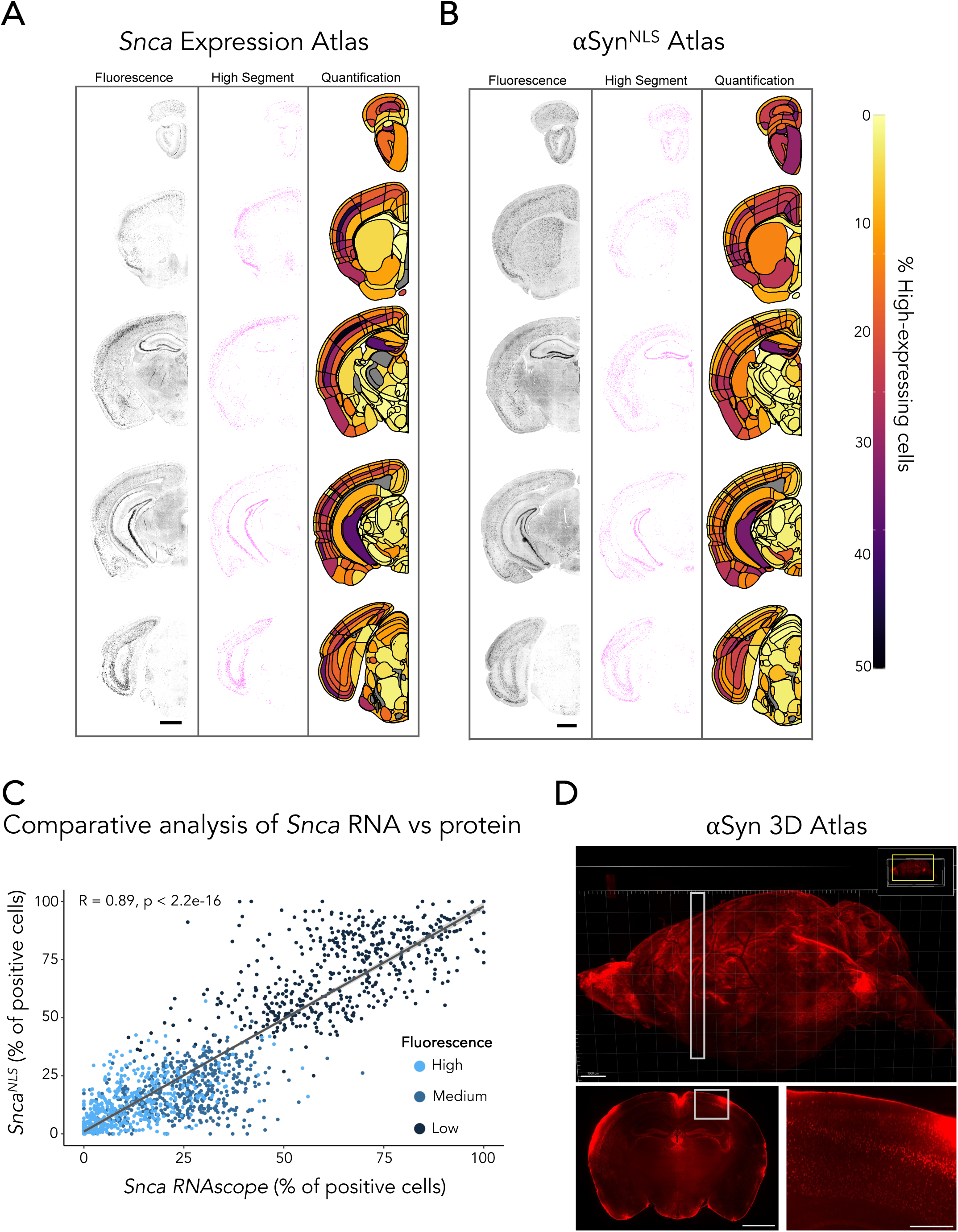
A brain-wide atlas of αSyn topography. A and B) RNAScope performed on a wild-type mouse to measure *Snca* expression (A) and immunofluorescent staining performed on a *Snca^NLS^* mouse brain measuring αSyn protein density (B) throughout the brain. Staining (left), segmentation (middle, high threshold), and heatmap (right) show brain region-specific expression patterns of *Snca* or αSyn distribution, respectively. Immunofluorescent staining performed on a *Snca^NLS^* mouse brain measuring αSyn protein density C) A comparative analysis of A and B indicating correlation of topography between *Snca* RNAScope and αSyn immunofluorescence for different intensity cut-offs. Dots represent class prevalence from individual regions. The solid line represents the line of best fit, and the shaded ribbons represent the 95% prediction intervals. The R and p values for the Pearson correlation between vulnerability and gene expression are noted on the plots. D) Representative images of whole brain Flag epitope staining and imaging provide an encompassing view of αSyn topography. White boxes indicate insets. Scale bars: 1,000 µm (A, B, D top, D bottom left), 250 μm (D bottom right).

Second, we performed serial sectioning and diaminobenzidine (DAB) staining for αSyn at defined coordinates, then quantified the relative density of αSyn+ cells throughout the brain. The DAB-stained tissue from both coronal (Supp **Fig 2A**) and sagittal (Supp **Fig 2B**) sections were analyzed using ilastik, an image classification and segmentation tool that uses machine learning to delineate αSyn+ cells^27^. Next, the αSyn+ cell density was averaged against the hematoxylin+ cell density to obtain the relative αSyn+ cell density per brain region (Supp **Fig 2C**).

**Figure 2:**
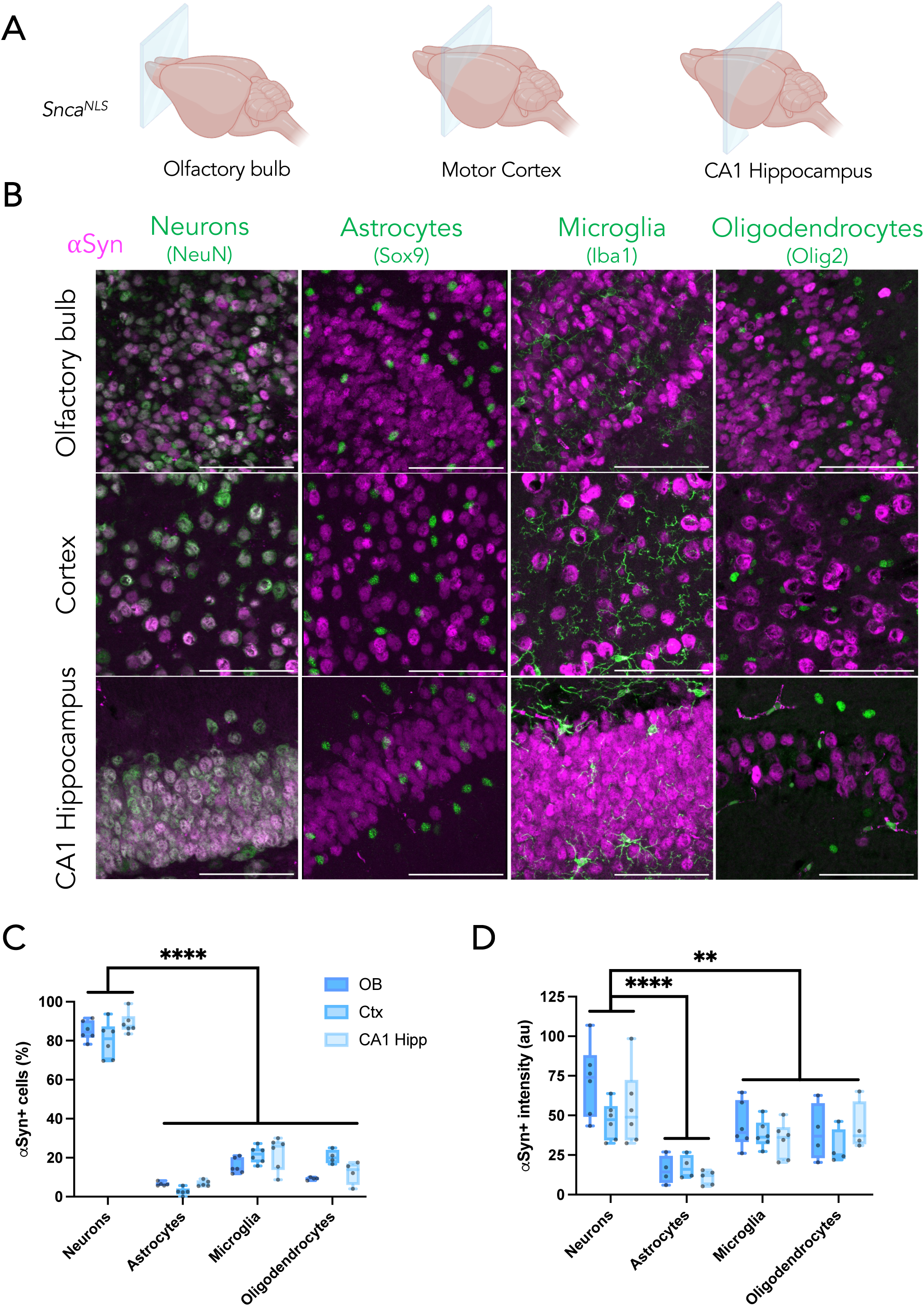
αSyn is predominantly expressed in neurons. A) Coronal planes chosen to assess different brain regions in *Snca^NLS^* mice. B) Merged micrographs from the olfactory bulb (upper), motor cortex (middle), and CA1 hippocampus (bottom) staining for neurons (far left), astrocytes (middle left), microglia (middle right), and oligodendrocytes (right). From these, αSyn density (C) and intensity (D) were quantified. Two-way ANOVA with Tukey’s post hoc, **, **** denote p<0.01 and p<0.0001, respectfully. Scale bars: 75 μm.

Third, to better understand the regional density of αSyn+ cells based on their expression levels, we generated a coronal atlas from IF-stained brain tissue at defined coordinates (Supp **Fig 1B**). This tissue was analyzed using an adapted version of the QUINT workflow^28^, which uses QuPath, QuickNII^29^, VisuAlign^30^, and Nutil^31^ to quantify cell classes in brains registered to the Allen Brain Atlas CCFv3 (**Fig 1B**). These three methods were all done in tandem and used to correlate RNA and protein topography (**Fig 1C**). In all methods tested, we observe largely concordant results suggesting that *Snca* RNA broadly correlates with αSyn protein.

Fourth, we performed whole-brain staining for αSyn, using the Flag epitope present in *Snca^NLS^* mice, coupled with light sheet microscopy to generate 3D renderings of αSyn topography throughout an intact brain (**Fig 1D**). Using this, we further explored the αSyn+ density and intensity from an intact brain, which largely agreed with our other methods (Supp **Fig 2D**). In all methods, we noted high αSyn expression throughout the cortex, especially cortical layers 4 and 5, with lower expression generally in layers 2/3 and 6. We also note high αSyn density and intensity in the hippocampus, and in the olfactory bulb, including the mitral, granule, and glomerular cell layers. αSyn expression was notably low in most thalamic and mesencephalic nuclei, except for the substantia nigra pars compacta (SNc).

### Neurons express variable αSyn levels

Pathological αSyn accumulates in multiple cell types, including monoaminergic neurons, glutamatergic neurons, and oligodendrocytes^32–34^. One hypothesized reason for this is that susceptible cells harbour more αSyn, thus rendering them more vulnerable to the aggregation process. The most common αSynucleinopathy, PD, is characterized by aggregates within neurons; however, a rarer αSynucleinopathy, MSA, is characterized by oligodendroglial cytoplasmic inclusions^2,4^. We sought to characterize αSyn-expressing cell types in brain regions with high αSyn+ cell density, namely the olfactory bulb, motor cortex, and hippocampus (**Fig 2A**). We co-stained *Snca^NLS^* mice for αSyn with cell-type markers for neurons (NeuN), astrocytes (Sox9), microglia (Iba1), and oligodendrocytes (Olig2; **Fig 2B**). Congruent with previous studies, αSyn is a primarily neuronal protein, where ∼80% of neurons in these brain regions consistently co-stain for αSyn, whereas only 10-20% of astrocytes, microglia, and oligodendrocytes express αSyn to a modest degree (**Fig 2C**). Neurons also express significantly more αSyn than all other cell types tested. Among the other cell types, microglia and oligodendrocytes have similar intensities of αSyn between one another, both generally higher than that of astrocytes (**Fig 2D**).

### Monoaminergic cells throughout the brain broadly and consistently express αSyn

Since PD is characterized by selective dopaminergic neurodegeneration – and more broadly, monoaminergic susceptibility^35^– we explored the density and intensity of αSyn protein in monoaminergic (TH+) cells of the ventral tegmental area (VTA), substantia nigra (SNc), and locus coeruleus (LC; **Fig 3A**). In PD, the SNc and LC have significant dopaminergic neurodegeneration, but the VTA is relatively spared^35^.Interestingly, we found that 100% of TH+ neurons in all three brain regions co-stain for αSyn (**Fig 3B, D**). Another brain region that is of importance in αSyn spreading, and PD degeneration is the dorsal motor nucleus of the vagus nerve (DMX). When staining for cholinergic neurons, we also found that 100% of ChAT+ cells of the DMX show αSyn+ staining (**Fig 3C, D**). The DMX-adjacent nucleus ambiguous (AMB), which is not known to accumulate αSyn pathology in PD^35^, was also positive for αSyn (**Fig 3E**). Intriguingly, we identified other ChAT+ cells, including the lateral reticular nucleus (LRN), that were devoid of αSyn+ staining (**Fig 3F**), suggesting that αSyn expression is not part of the essential ChAT+ neuron gene expression pattern. Despite this, the LRN has been shown to harbour significant αSyn pathology in disease^36^.

**Figure 3:**
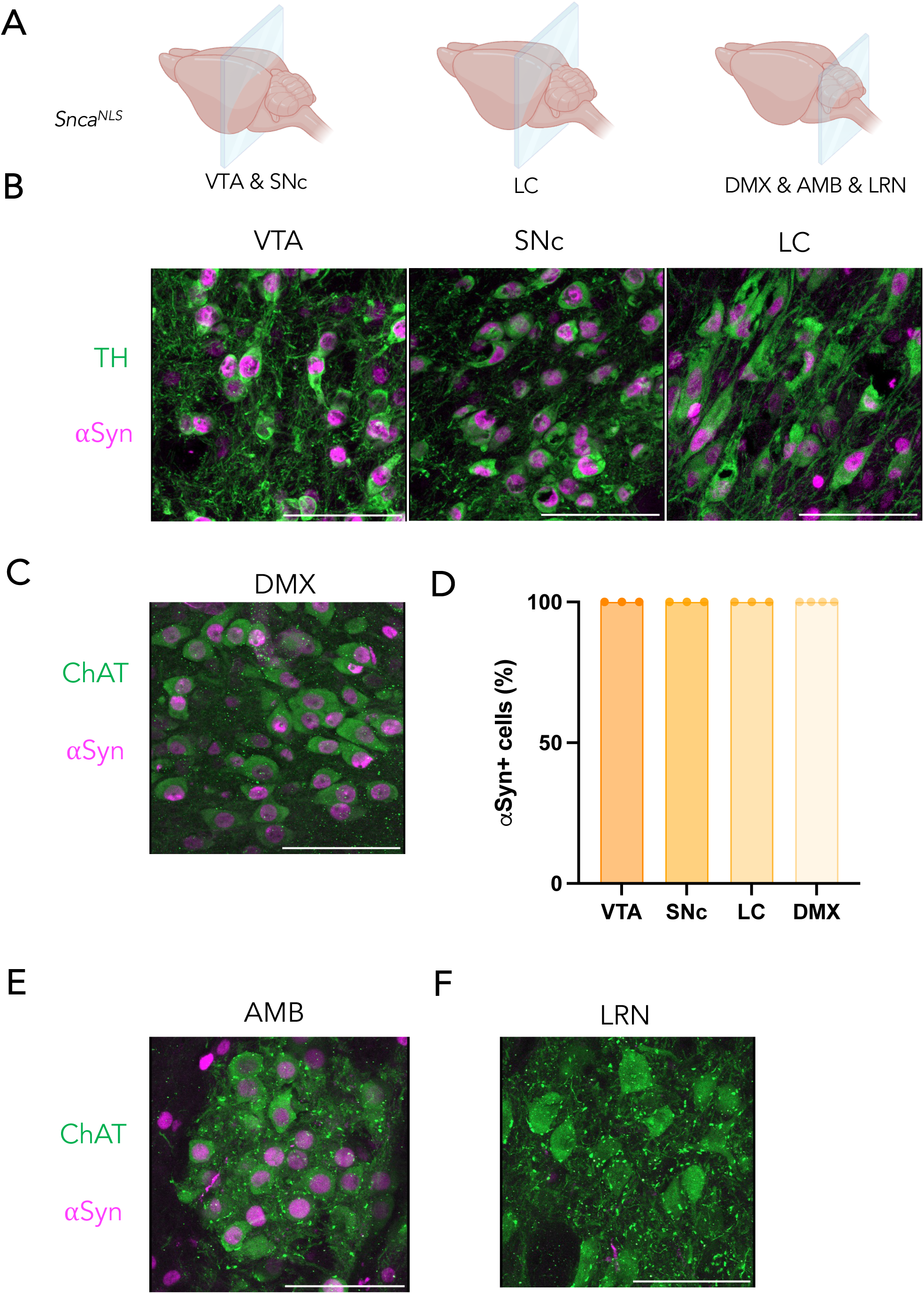
αSyn is highly expressed in catecholaminergic and cholinergic cell types that are vulnerable in PD. A) Coronal planes chosen to assess different brain regions in *Snca^NLS^*mice. B) TH+ cells of the ventral tegmental area (VTA, left), substantia nigra *pars compacta* (SNc, middle), and locus coeruleus (LC, right) as well as (C) ChAT+ cells of the dorsal motor nucleus of the vagus nerve (DMX) show 100% overlap with αSyn+ cells, quantified in (D). ChAT+ cells of the nucleus ambiguous (AMB, E) and lateral reticular nucleus (LRN, F) express high, and low αSyn expression, respectively. Scale bars: 75 μm.

### Spinal cord neurons exhibit heterogeneous αSyn expression patterns

The spinal cord is not routinely studied in PD, but has gained recognition for its position as a conduit between the gut and brain and its potential role in the gut-to-brain transmission of αSyn pathology^34,37–39^. Therefore, we next assessed αSyn staining patterns in the spinal cord of *Snca^NLS^* mice. We analyzed sections from the cervical, thoracic and lumbar spinal cord (**Fig 4A, Supp** **Fig 3A, 4A**). Within these regions, we analyzed the dorsal horn, ventral horn, and ventral white matter. As in the brain, αSyn in the spinal cord is primarily neuronal; however, its localization is not homogenous (**Fig 4B, Supp** **Fig 3B, 4B**). αSyn is expressed primarily in the dorsal horn, more so in upper laminae, with αSyn+ cells only appearing in the ventral horn more caudally (**Fig 4C, Supp** **Fig 3C**). Additionally, αSyn intensity varies along the length of the cord, with the brightest, most concentrated αSyn+ cells existing in the thoracic region (**Fig 4D, Supp** **Fig 3D**). Nearly half of all NeuN+ cells of the thoracic and lumbar dorsal horn co-label with αSyn, a fraction significantly smaller than the neurons of the brain (>80%). Similar to the brain, however, astrocytes, microglia, and oligodendrocytes have very few cells that co-label with αSyn. Additionally, the intensity of αSyn in the dorsal horn is higher than that of the ventral horn and ventral white matter, and is more consistent across cell types. These trends are all consistent in the cervical spinal cord (**Supp** **Fig 4**).

**Figure 4:**
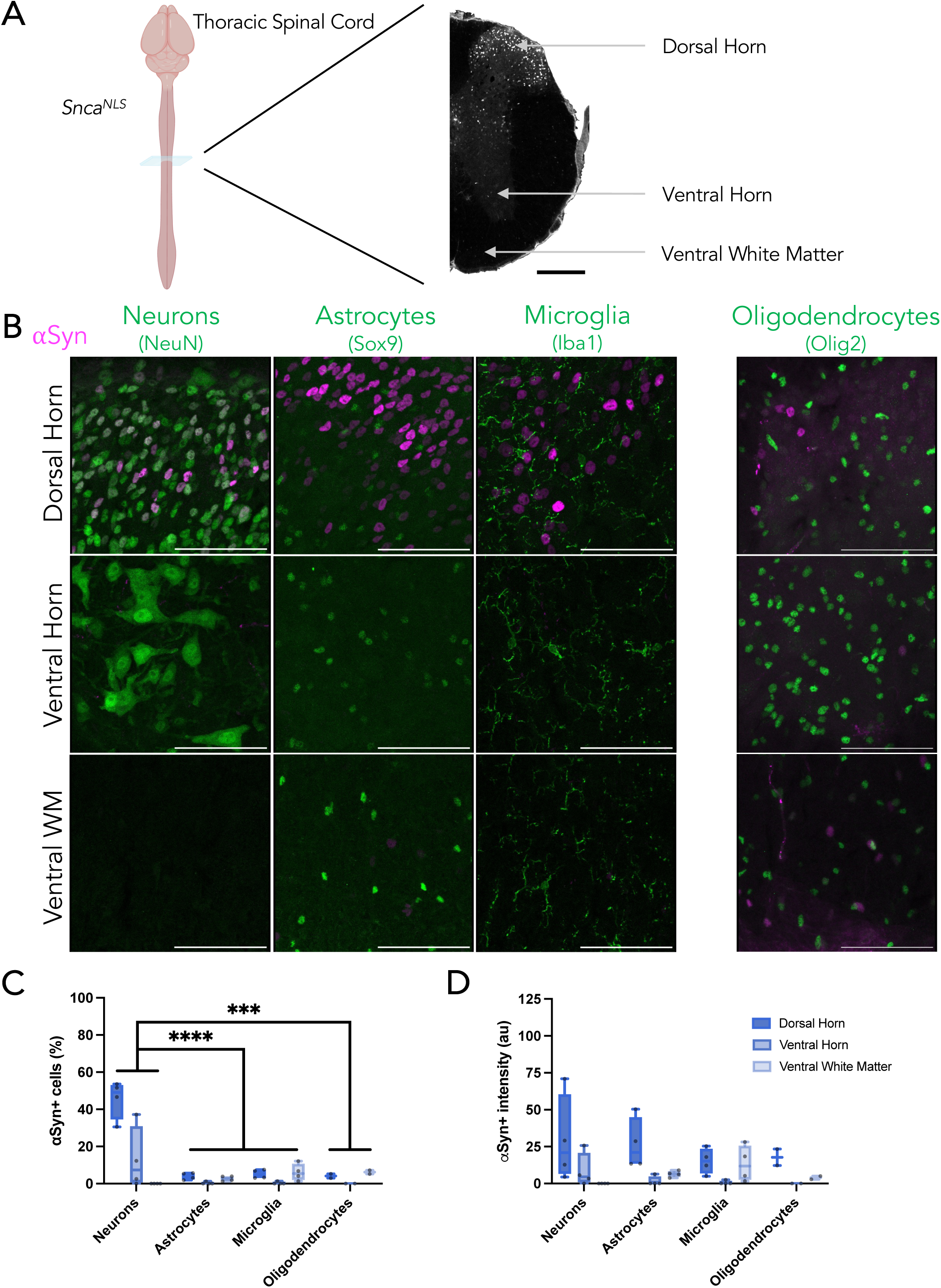
Diverse αSyn topography within the mouse spinal cord. A) Coronal plane chosen to assess different regions of the thoracic spinal cord in *Snca^NLS^* mice (left) with annotations for the dorsal and ventral horns and ventral white matter (right) from a section stained for αSyn. B) Merged micrographs from the dorsal horn (upper), ventral horn (middle), and ventral white matter (bottom) staining for neurons (left), astrocytes (middle left), microglia (middle right), and oligodendrocytes (right; different staining paradigm). From these, αSyn density (C) and intensity (D) were quantified. Two-way ANOVA with Tukey’s post hoc, ***, **** denotes p<0.001 and p<0.0001, respectively. Scale bars: 1000 μm (A), 75μm (B).

To further investigate the neuronal localization of αSyn in the spinal cord, we co-stained cervical, thoracic, and lumbar sections with ChAT, a marker of motor neurons, and Pax2, a marker of GABAergic neurons. In both cases, we found that αSyn colocalizes with these markers primarily within lamina X surrounding the central canal (**Supp** **Fig 5**). Since αSyn+ neurons represent only 50% of the entire spinal cord neuron population (NeuN+; compared to >80% in the brain regions tested, **Fig. 2**), we turned to the harmonized atlas of single cell sequencing data to validate these findings^40,41^. Consistent with our imaging, we found that *<u>Snca</u>* was enriched in neurons (**Supp** **Fig 6A, A’, A”**), specifically excitatory clusters 18 and 19 within the Sox5 family (distinguished by *Nmu* and *Tac2* expression, respectively), and inhibitory clusters 9-11 within the Pdyn family. Similarly, the harmonized atlas also illustrates the lack of overlap between *Snca* and motor neurons (**Supp** **Fig 6A”**). Additionally, scRNA-Seq datasets from the mouse brain^42,43^ and colon^44,45^ show similar congruency with our atlas, illustrating the abundance of neuronal αSyn with specific sub-neuronal clusters exhibiting higher αSyn expression (**Supp** **Fig 6B-C”**). These data demonstrate that αSyn exhibits a neuron subtype-specificity in the spinal cord.

**Figure 5:**
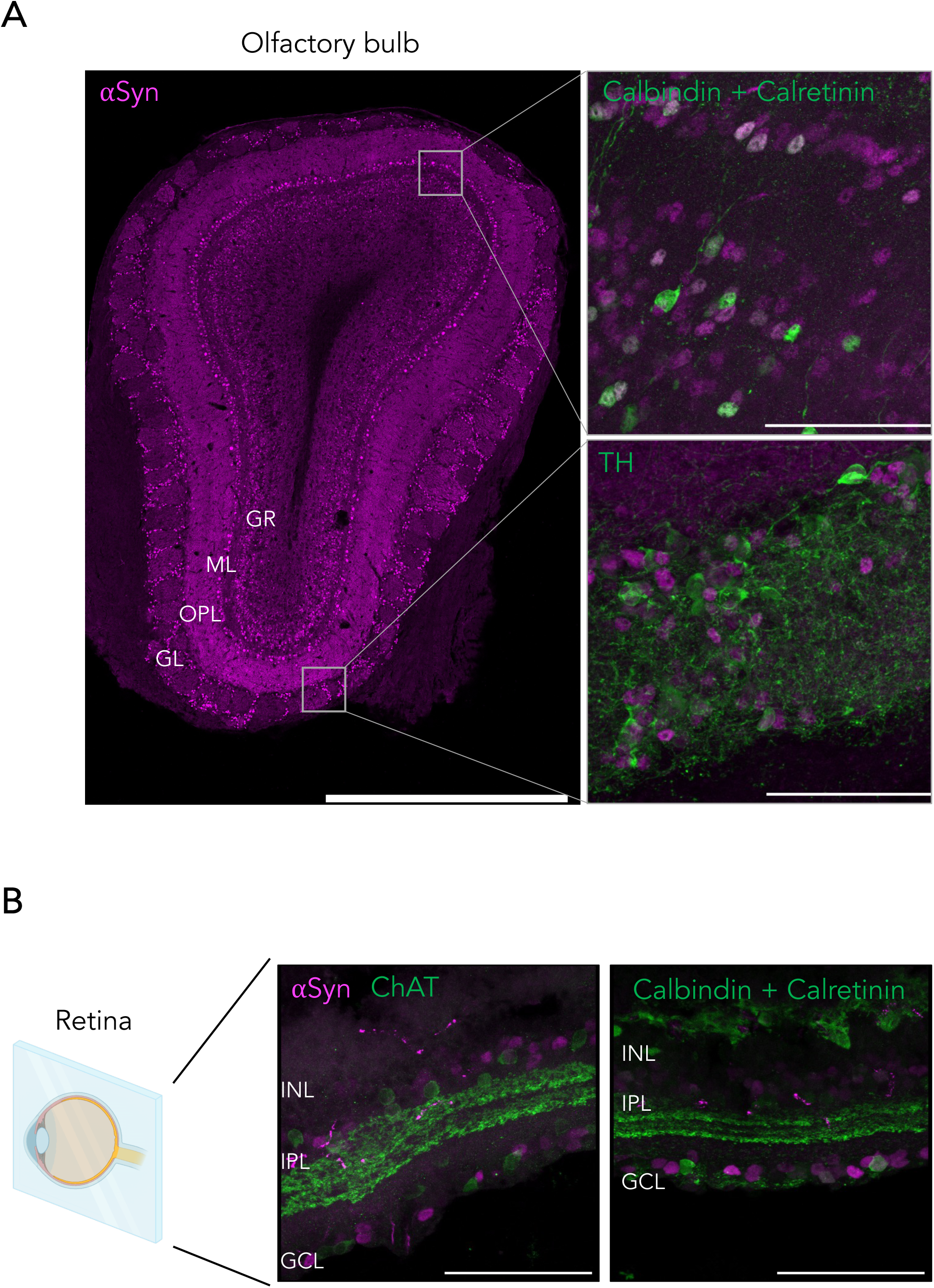
Diverse αSyn staining patterns within the mouse olfactory bulb and retina. A) Tiled immunofluorescent image of an olfactory bulb (left) shows region-specific topography of αSyn, specifically colocalizing with calbindin/calretinin in the mitral layer (upper right) and TH in the glomerular layer (lower right). B) Cross section of *Snca^NLS^* retina (left) co-staining for ChAT (middle) or calbindin/calretinin (right) show distinct staining patterns in the ganglionic cell layer (GCL) and inner plexiform layer (IPL). Scale bars: 1,000 μm (A left), 75 μm (A right, B).

**Figure 6:**
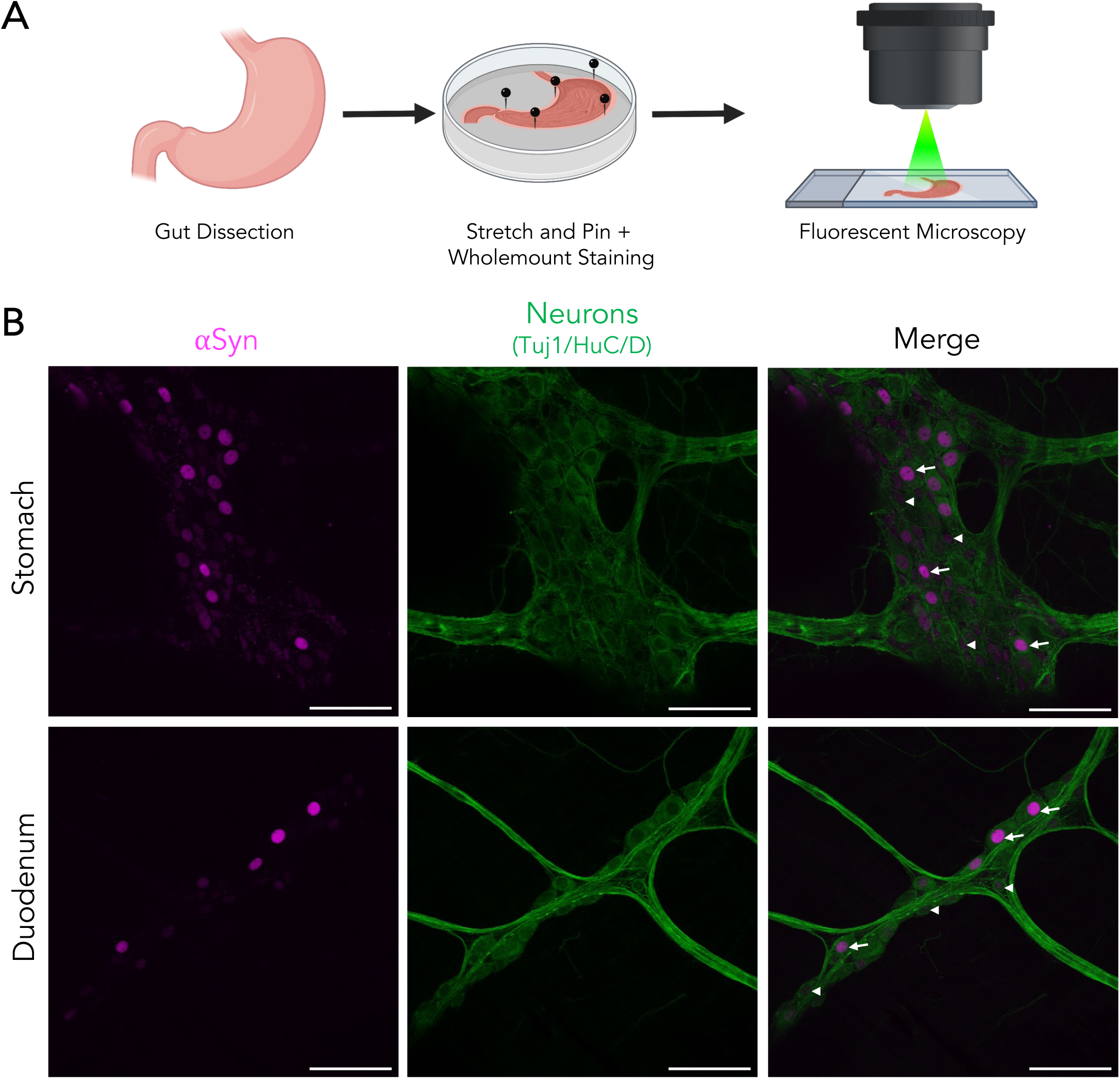
Varying intensities of αSyn expression throughout the gut. A) *Snca^NLS^* gut tissue was flushed and stained whole-mount prior to immunofluorescent staining and imaging. B) αSyn shows 100% colocalization with neuronal markers in the stomach (upper) and duodenum (lower). Arrow heads denote low αSyn, arrows denote high αSyn intensity. Scale bars: 75 μm.

### Olfactory bulb and retina show distinct subpopulations of cells with high αSyn density and intensity

The “brain-first” model of PD suggests that αSyn pathological spread begins in the brain or olfactory bulb before moving into the periphery^46–48^. Additional *in vivo* work has implicated the olfactory bulb as a main hub for αSyn pathology, specifically in the mitral cells^49^. We therefore investigated the olfactory bulb in more detail in *Snca^NLS^* mice. We found prominent αSyn staining in the mitral and glomerular regions of the olfactory bulb, where we found that αSyn colocalizes with all TH+ glomerular cells and all calbindin/calretinin+ cells in the granular and mitral layers (**Fig 5A**).

In conjunction with the olfactory bulb, recent studies have explored the retina as a site for αSyn pathology genesis and spread in the brain-first model. One group found intravitreal injection of αSyn PFFs led to αSyn pathology, retinal degeneration, and dopaminergic deficits^50^. Other groups show increased αSyn pathology in the retina and optic nerve, with retinal neurodegeneration in PD and other αSynucleinopathy patients^51,52^. These studies typically report pathology and neurodegeneration in the ganglionic cell layer and inner plexiform layer. Incidentally, these sites of pathology are where we noted most αSyn+ cells. Upon co-staining with cell-specific markers, we noted αSyn does not colocalize with ChAT+ neurons, but with calbindin/calretinin+ neurons in the ganglion and inner plexiform layers (**Fig 5B**). We also noted higher αSyn intensity in the ganglion cell layer than in the inner plexiform layer.

### All enteric neurons of the stomach and duodenum have αSyn protein of varying intensity

A counterpart to the brain-first hypothesis is the “body-first” model, where αSyn pathology begins in the gut and travels retrogradely to the brain^46,48^. Previous studies have injected αSyn PFFs into the gut and observed spreading into the brain^53–55^; however, the cells responsible for this spreading remain unknown. We therefore characterized αSyn protein expression in the gut, as a potential site for the initiation of propagation of αSyn pathology. Since we previously noted that αSyn primarily resides in neurons, we focused on the enteric neurons. We performed whole-mount staining of the stomach and duodenum of *Snca^NLS^*mice then co-stained for αSyn and Tuj1/HuC/D to mark enteric neurons (**Fig 6A**). This whole-mount preparation preserves both enteric neurons and longitudinal muscle. Here, we find αSyn exclusively within enteric neurons of both the stomach and duodenum with considerable variability in the relative amount of αSyn between neurons (**Fig 6B**). This finding is consistent across the stomach and duodenum and is in line with recent enteric nervous system single-cell sequencing findings (Supp Fig 6C, C’, C”)^45,56^.

## Discussion

In this study, we used our *Snca^NLS^* mouse model to characterize the topography of αSyn systematically throughout the brain and in other PD-relevant tissues. We found that αSyn is expressed predominantly in neurons and that, while it is expressed in some of the non-neuronal cells tested, it is at much lower levels. To our surprise, cholinergic neurons express αSyn to varying degrees, with some expressing high levels of αSyn (e.g. DMX) while others are virtually devoid of αSyn (e.g. motor neurons of the spinal cord). It was interesting to note that olfactory neurons (particularly TH-positive, periglomerular cells) were highly enriched for αSyn whereas retinal ganglion cells and amacrine cells expressed very low levels of αSyn, which bordered on the lower limit of detection. This is consistent with recent findings that suggest that the olfactory bulb is likely a richer source of seed-competent αSyn than the retina^50,57–59^.

Astrocytes, microglia, and oligodendrocytes all expressed αSyn but at very low levels. Some populations of Olig2+ cells, however, expressed a considerable amount of αSyn. These are likely oligodendrocyte precursor cells (OPCs) which have previously been reported to express a relatively high level of αSyn^60,33,61–64^. The role that αSyn plays within these cells remains enigmatic, but higher native expression in OPCs, in concert with genetic and/or environmental insults, may predispose a cell to future pathology in diseases like MSA, where αSyn pathology principally occurs in oligodendrocytes.

Myenteric neurons express a considerable amount of αSyn. Interestingly, this level is highly varied between neuronal subtypes. This finding is consistent with a recent single cell transcriptomic dataset that highlights an enrichment of *Snca* in enteric neuron classes (ENCs) 5, 9, 10 and 11^56^. Interestingly, ENC11 is defined by a high expression of TH and Calb2, markers that cluster with high αSyn expressing cells like dopamine neurons.

The development of this topographical atlas of αSyn expression sets the stage for future studies exploring the impact of neuronal αSyn on subsequent vulnerability to developing Lewy pathology. First, this atlas provides a newfound level of insight in healthy, young animals. Future work, looking at the impact of PD-relevant risk factors such as age, toxin or genetic exposure, or viral insults will shed important light on modes of native αSyn regulation. Second, it remains unclear whether the native abundance of αSyn affects the susceptibility of a cell to neurodegeneration. Seeding-based experiments suggest that cells with higher αSyn are more vulnerable to pathology^65,66^, while human imaging studies have suggested that regional atrophy in PD is related to anatomical connectivity^67^. It is likely that these two factors (anatomical connectivity and gene expression) work in concert to make certain neurons and brain regions vulnerable to developing Lewy pathology^68^. Future work looking to selectively target αSyn-rich versus αSyn-poor circuits will provide much-needed understanding of the role of endogenous αSyn dosage and cellular vulnerability. Lastly, given the increased focus on αSyn-lowering therapeutics (e.g. immunotherapy, antisense oligonucleotides, aggregation inhibitors, viral approaches), this atlas will provide insight into why certain cell types may be more resistant to these translational modalities, providing ways to better target therapeutics and circumvent associated pitfalls.

## Methods

### Mouse Husbandry

All mice were housed with up to 5 mice per cage on a 12-hour light-dark cycle. Mice were fed *ad libitum* and all husbandry was performed by the uOttawa Animal Care and Veterinary Services staff. All animal work was performed under the breeding protocols (CMMb-3654 and CMMb-3904) approved by the uOttawa Animal Care Committee. The *Snca^NLS^* mouse line used in this study is available through the Jackson Laboratory (Jax Stock No. 036763). All housing and procedures at Van Andel Institute were performed according to the NIH Guide for the Care and Use of Experimental Animals and approved by the Van Andel Institute Institutional Animal Care and Use Committee (IACUC). C57BL/6J mice were purchased from the Jackson Laboratory (000664; RRID:IMSR_JAX:000664).

### Tissue collection

Mice were sedated with an intraperitoneal injection of 120mg/kg Euthanyl (DIN00141704). Next, they were transcardially perfused with 10mL 1X phosphate buffered saline (PBS) + 10U/mL heparin (Millipore Sigma, H3393-50KU) followed by 10mL 4% paraformaldehyde (PFA). Brains were extracted and incubated in 4% PFA overnight at 4°C with gentle rocking. The brains were then washed twice with 1X PBS (**<u>dx.doi.org/10.17504/protocols.io.b5swq6fe</u>**). Brains used for RNAscope and immunofluorescence mapping were processed into paraffin via sequential dehydration and perfusion with paraffin under vacuum (70% ethanol for 1 hour, 80% ethanol for 1 hour, 2 times 95% ethanol for 1 hour, 3 times 100% ethanol for 1 hour, 2 times xylene for 30 minutes, paraffin for 30 minutes at 60°C, paraffin for 45 minutes at 60°C). Brains were then embedded in paraffin blocks, cut into 6 µm sections and mounted on glass slides. Subsequent steps are protocol-dependent and outlined in downstream methods. See **Supplementary Table 1** for a comprehensive list of all antibodies/kits used in this study.

### *Snca* Expression, αSyn^NLS^, and αSyn DAB Atlas

#### In situ hybridization

In situ hybridization was performed with RNAscope® Multiplex Fluorescent Reagent Kit v2 (ACD, Cat #323270) using recommended conditions and supplied reagents. Paraffin-embedded tissue was freshly sectioned and dried. When it was not going to be used immediately, slides were vacuum-sealed and stored at 4°C. Slides were baked in a dry oven for 1 hour at 60°C and used within one week. Slides were de-paraffinized with 2 sequential 5-minute washes in xylenes, followed by 2 washes in 100% ethanol for 2 minutes. Slides were then dried for 5 minutes at 60°C. Slides were treated with hydrogen peroxide for 10 minutes at room temperature and washed two times with distilled water. Target retrieval was performed in target retrieval reagents in the BioGenex EZ-Retriever System for 15 minutes at 99°C. Slides were then washed with distilled water for 15 seconds and transferred to 100% ethanol for 3 minutes before being dried for 5 minutes at 60°C.

Slides were incubated in protease plus in a humidified tray in a hybridization oven (Boekel Scientific 240200) for 30 minutes at 40°C. Slides were washed 2 times with distilled water. RNAscope® probes were added to slides and incubated for 2 hours at 40°C. The following probe was used: *Snca* (RNAscope probe, Mm-Snca-C1, 313281). Slides were washed twice for 2 minutes with wash buffer and incubated in Amp 1 for 30 minutes at 40°C. The wash and Amp incubation was repeated for Amp 2 and Amp 3, except Amp 3 was only incubated for 15 minutes. Slides were washed twice for 2 minutes with wash buffer and incubated in HRP-C1 for 15 minutes at 40°C. Slides were washed twice for 2 minutes with wash buffer and incubated in Opal 520 (Perkin Elmer FP1487A) for 30 minutes at 40°C, washed twice for 2 minutes with wash buffer, incubated in HRP blocker for 15 minutes at 40°C, and washed twice for 2 minutes with wash buffer. Slides were mounted with coverglass in ProLong gold with DAPI (Invitrogen, Cat#P36931) and imaged at 20x magnification on a Zeiss AxioScan 7 microscope.

#### Immunofluorescence

Slides were de-paraffinized with 2 sequential 5-minute washes in xylenes, followed by 1-minute washes in a descending series of ethanols: 100%, 100%, 95%, 80%, 70%. Slides were then incubated in deionized water for one minute prior and transferred to the BioGenex EZ-Retriever System where they were incubated in antigen unmasking solution (Vector Laboratories; Cat# H-3300) and microwaved for 15 minutes at 95°C. Slides were allowed to cool for 20 minutes at room temperature and washed in running tap water for 10 minutes. Slides were washed for 5 minutes in 0.1 M Tris, then blocked in 0.1 M Tris/2% fetal bovine serum (FBS) for 2 hours. Slides were incubated in primary antibody in 0.1 M Tris/2% FBS in a humidified chamber overnight at 4°C.

Primary antibodies were rinsed off with 0.1 M tris and incubated in 0.1 M Tris/2% FBS for 5 minutes. Slides were then incubated in the dark for 3 hours at room temperature with secondary antibodies in 0.1 M Tris/2% FBS. Slides were rinsed three times for 10 minutes in 0.1 M Tris in the dark, then mounted with coverglass in ProLong gold with DAPI (Invitrogen, Cat#P36931). Fluorescent slides were imaged at 20X magnification on a Zeiss AxioScan 7 microscope.

#### Cell detection and classification

Our study included 99 sections from 4 *Snca^NLS^* mice for immunofluorescence and 49 sections from 4 C57BL/6J mice for *in situ* hybridization. Stained slides scanned on a Zeiss AxioScan 7 at 20x magnification were imported into QuPath v0.2.3 or newer for analysis. Individual cells were identified using the Cell Detection feature that allows detection based on DAPI stain intensity. Cell detection parameters such as background radius, sigma, and threshold were adjusted to optimize cell detection across all brain sections.

After cell detection, cell classification was completed in QuPath using the object classification feature. QuPath was trained on a subset of annotations to distinguish between different cell types based on signal intensity. To train the classifier, cells were detected by DAPI then classified according to the nuclear expression intensity of Syn1 staining or whole-cell intensity of *Snca* staining. Thresholds for the intensity classifications were calculated by averaging the mean intensity of all cells. Three classes were then generated: 1) Low: one standard deviation or less below the mean cellular intensity; 2) Medium: one standard deviation below or above the mean cellular intensity; 3) High: one standard deviation or more above the mean cellular intensity. This classifier was then applied to all brain sections. Each class was assigned a different colour and exported as .png files for subsequent analysis.

#### Mouse Brain Registration

Images were registered to the Allen Brain Atlas CCFv3 using a modified version of the QUINT workflow^21^. An RGB image of each section was exported from QuPath as a .png file, downsampled by a factor of 12, to use for spatial registration in QuickNII^22^. A segmentation image was created by exporting a colour-coded image of classified cells by category on a white background to use as the segmentation input in Nutil. Brain images were uploaded in Filebuilder and saved as an XML file to be compatible with QuickNII. Following the spatial registration of the mouse brain sections to the Allen Mouse Brain Atlas CCFv3 in QuickNII, a JSON file was saved for use in (VisuAlign, RRID:SCR_017978). Brain sections were imported into VisuAlign to perform non-linear warp transformations and align all brain regions. Anchor points were generated in the atlas overlay and moved to the corresponding location on the brain section. Final alignments were exported as .flat and .png files for use in Nutil^78^.

Nutil was used for the quantification and spatial analysis of the identified cell types in specific regions of the mouse brain. Individual classes were identified for quantification via their HTML colour code assigned in QuPath. Nutil generated object counts from each individual classification within each region of the Allen Mouse Brain Atlas using the registration from QuickNII and VisuAlign. The percentage of each class was calculated for each region and used for subsequent plotting and analysis. Anatomical heatmaps were generated using custom R code (https://github.com/vari-bbc/Mouse_Brain_Heatmap).

#### DAB staining

Cryosectioned tissue were incubated in 0.9% hydrogen peroxide (Millipore Sigma, 216763-500ML) for 10 minutes at room temperature. The following diaminobenzidine staining was performed using the Vectastain Elite ABC HRP Mouse Kit (Vector Laboratories, PK-6102). Briefly, tissue were incubated in blocking buffer (1.5% triton X-100 + 5% Vectastain Horse Serum in 1X PBS) for 1 hour at room temperature following by incubation in primary antibody solution overnight at 4°C. Next, tissue was incubated in secondary antibody diluted in blocking buffer for 1 hour, then ‘A+B’ solution (mixed 30 minutes prior to use) for 1 hour at room temperature. Tissue was then exposed to diaminobenzidine for 2 minutes using the DAB Peroxidase HRP Substrate Kit (Vector Laboratories, SK-4100) then washed with 1X PBS. Tissue was dehydrated in 50%, 70%, 95%, then 100% ethanol for 1 minute each, then 100% xylenes for 5 minutes. Finally, tissue was coverslipped with Permount mounting medium (Fisher Scientific, SP15-100) and #1.5 coverslips. Stained slides were imaged at 20X magnification using an Axio Scan Z1 Slide Scanner and exported as czi files for subsequent analysis (dx.doi.org/10.17504/protocols.io.b5s9q6h6).

#### Hematoxylin staining

Cryosectioned tissues were stained using the H&E Staining Kit (Abcam, ab245880) following kit instructions. Briefly, free-floating sections were incubated in hematoxylin for 5 minutes then washed with 2 changes of water, blueing reagent for 15 seconds, then washed in water again. Tissue was then mounted on a superfrost plus slide and dehydrated in 50%, 70%, 95%, then 100% ethanol for 1 minute each then 100% xylenes for 5 minutes before coverslipping with Permount mounting medium (Fisher Scientific, SP15-100) and #1.5 coverslips. Stained slides were imaged at 20X magnification using the Axio Scan Z1 Slide Scanner and exported as .czi files for subsequent analysis.

#### αSyn DAB Atlas quantification

All tissue slices were annotated to specific Allen Brain Atlas coordinates then converted to tiff files using Fiji ImageJ (version 2.3.0/1.53q). The ‘pixel classification’ pipeline from ilastik software (version 1.3.2) was used for to delineate positive cells and three images were used in training the machine learning software. Segmented images were opened in Fiji and brain regions were manually outlined and added to an ROI manager. The ‘analyze particles’ function was used to count positive cells greater than 5 pixels in size, corresponding to 1 positive cell. The number of αSyn+ cells was divided by the number of hematoxylin positive cells and multiplied by 100 to give then relative αSyn+ cell density.

#### Whole brain clearing and imaging

Whole *Snca^NLS^* mouse brains were processed using the SHIELD protocol by LifeCanvas Technologies^69^. Samples were cleared for 7 days with Clear+ delipidation buffer then immunolabeled using SmartLabel. Each sample was labelled with 5μg mouse anti-NeuN (RRID:AB_2572267) and 4μg goat anti-Flag (RRID:AB_299216) followed by fluorescent secondary antibodies. Samples were incubated in EasyIndex for a refractive index of 1.52 and imaged at 3.6X using SmartSPIM microscopy. Images were tile-corrected, de-striped, and registered to the Allen Brain Atlas (https://portal.brain-mapping.org). The NeuN channel was registered to the 8-20 atlas-aligned reference samples using successive rigid, affine, and b-spline warping (SimpleElastix: https://simpleelastix.github.io). Average alignment to the atlas was generated across all intermediate reference sample alignments to serve as the final atlas alignment value per sample. Fluorescent measurements from the acquired images were projected onto the Allen Brain Atlas to quantify the total fluorescence intensity per region defined by the Allen Brain Atlas. These values were then divided by the volume of the corresponding regional volume to calculate the intensity per voxel measurements. See **Supplementary Table 2** for raw output files from LifeCanvas. Data was plotted using Prism (version 9.5.1).

### Cell-specific staining and quantification

#### Organ isolation

Following intracardial perfusion (see tissue collection method above), brains, spinal cords, and retina from *Snca^NLS^* mice were extracted and incubated in 4% PFA for 72 hours at 4°C with gentle rocking. Brain and spinal cords were next dehydrated in a 3-step sucrose sequence with 10, 20, and 30% sucrose for 24 hours each. Finally, brains and spinal cord were flash frozen for 1 minute at -40°C isopentane, suspended in OCT, then cryosectioned at 40μm. Eyes were incubated in 70% ethanol following fixation, then retinas were isolated, suspended in OCT, and cryosectioned at 20μm (dx.doi.org/10.17504/protocols.io.b5swq6fe).

#### Immunofluorescent Staining

Cryosectioned, free-floating tissue sections were incubated in blocking buffer (0.5% Triton X-100 (Sigma-Aldrich, T8787-100ML) + 10% normal horse serum (Sigma-Aldrich, H0146-5ML) in 1X PBS) for 1 hour at room temperature. Next, sections were incubated in 1° antibody overnight at 4°C then fluorescently conjugated 2° antibody for 1 hour at room temperature. Free-floating sections were mounted on SuperFrost+ slides then coverslipped with fluorescent mounting medium (Agilent, S302380-2) and #1.5 coverslips (dx.doi.org/10.17504/protocols.io.b5s5q6g6).

#### ChAT staining

Cryosectioned tissues were incubated in blocking buffer (0.1% Triton X-100 (Sigma-Aldrich, T8787-100ML) + 10% normal horse serum (Sigma-Aldrich, H0146-5ML) + 0.5% gelatin (Sigma-Aldrich, G1393-100ML) in 1X PBS) for 2 hours at room temperature. Next, sections were incubated in 1° antibody for 48 hours at 4°C then fluorescently conjugated 2° antibody for 2 hours at room temperature. Free floating sections were mounted on SuperFrost+ slides then coverslipped with fluorescent mounting medium (Agilent, S302380-2) and #1.5 coverslips (dx.doi.org/10.17504/protocols.io.rm7vzxbz4gx1/v1).

#### Brain quantification

Stained sections were imaged with a Zeiss LSM800 AxioObserver Z1 microscope at 20X with Z-stack. Images were then 3D projected and all channels were merged. Each αSyn+ cell nucleus that co-labels with the specific cell marker was circled and the intensity was measured in Fiji ImageJ (version 2.3.0/1.53q) with the ROI manager. The background signal was measured by circling 3 separate regions of αSyn-space and measuring the intensity with the ROI manager. To quantify, the background values were averaged and subtracted from each cell-specific intensity measurement. These intensity values were plotted in a nested graph using Prism (version 9.5.1) and analyzed using two-way ANOVA with Tukey’s post hoc analysis.

#### ENS immunofluorescence

Following euthanasia and perfusion of animals, stomach and duodenum were collected, washed, stretched, pinned flat on Sylgard-coated Petri dishes, and fixed overnight at 4°C in phosphate buffer saline solution (PBS) containing 4% (vol/vol) paraformaldehyde (cat# P6148, Sigma-Aldrich). Layers of tissue containing the myenteric plexus were separated by microdissection. Tissues were permeabilized at room temperature (RT) for 2h in a 10% fetal bovine serum (FBS)/PBS blocking buffer containing 0.5% Triton X-100 (cat# X100, Sigma-Aldrich), and incubated overnight at 4°C with the following primary antibodies diluted in the blocking buffer: rabbit monoclonal anti-alpha-synuclein. Following incubation with primary antibodies, tissues were washed with PBS and incubated for 1h at RT in secondary antibodies. Tissues were then washed with PBS and mounted with ProLong Gold antifade reagent (cat# P36931, Thermo Fisher Scientific). Confocal images were acquired using a Nikon A1plus-RSi confocal microscopy under a 40x objective and were analyzed with Image J software (dx.doi.org/ 10.17504/<u>protocols.io</u>.14egn3wxpl5d/v1).

## Author contributions

H.M.G., Z.A.F., M.X.H. and M.W.C.R. conceived the study; H.M.G., Z.A.F., L.S., A.P., K.K., and S.M.C. collected primary data. H.M.G., Z.A.F., L.S. and A.P., M.X.H. and M.W.C.R. analyzed data. H.M.G, M.X.H and M.W.C.R. wrote the paper.

## Acknowledgements

The authors thank Michelle Seguin and Meghan Heer for valuable insight into the early phases of this project and assistance with colony maintenance. This research was funded in part by Canadian Institutes of Health Research (PJT-169097, to M.W.C.R.); the Parkinson Canada New Investigator Award (2018-00016, to M.W.C.R.); Aligning Science Across Parkinson’s [ASAP-020625 to M.W.C.R., ASAP-020616 to M.X.H.] through the Michael J. Fox Foundation for Parkinson’s Research (MJFF); Canadian Institutes of Health Research Canada Graduate Scholarship-D (FBD-181526, to H.M.G.); Dave and Jill Hogg and Toth Family Parkinson Research Consortium Fellowships (Z.A.F.). For the purpose of open access, the author has applied a CC BY public copyright license to all Author Accepted Manuscripts arising from this submission. The authors thank the following Core facilities from the University of Ottawa for use of their facility, equipment, and expertise: Cell Biology and Imaging Acquisition Core (RRID: SCR_021845) and Louise Pelletier Histology Core (RRID: SCR_021737). Figures. 2A, 3A, 4A, 5B, 6A, Supplemental Figures 3A, and 4A were generated, in part, using Biorender.com. All histograms were generated with Prism 9.

## Conflicts of interest

None to declare.

**Supplemental Figure 1:** Brain-wide heatmaps of *Snca* expression and αSyn protein density. A) *Snca* intensity from RNAScope of a wild-type mouse binned into low (upper panel), medium (middle panel), and high intensity (lower panel) prior to generating the heatmaps. B) αSyn staining intensity of a *Snca^NLS^* mouse binned into low (upper panel), medium (middle panel), and high intensity (lower panel prior to generating the heatmaps.

**Supplemental Figure 2:** Brain-wide atlas of αSyn protein density and intensity. A) Coronal atlas of αSyn DAB staining (left) and ilastik segmentation (right). B) Sagittal atlas of αSyn ilastik segmentation following DAB staining. C) Quantification of αSyn density in targeted brain regions. D) Quantification of αSyn+ density and intensity from whole brain staining and imaging.

**Supplemental Figure 3:** αSyn density and intensity of the lumbar spinal cord. A) Coronal plane chosen to assess different regions of the lumbar spinal cord in *Snca^NLS^* mice (left) with annotations for the dorsal and ventral horns and ventral white matter (right) from a section stained for αSyn. B) Merged micrographs from the dorsal horn (upper), ventral horn (middle), and ventral white matter (bottom) staining for neurons (left), astrocytes (middle left), microglia (middle right), and oligodendrocytes (right; different staining paradigm). From these, αSyndensity (C) and intensity (D) were quantified. Two-way ANOVA with Tukey’s post hoc, **** denotes p<0.0001. Scale bars: 1,000 μm (A), 75μm (B).

**Supplemental Figure 4:** αSyn staining of the cervical spinal cord. A) Coronal plane chosen to assess different regions of the lumbar spinal cord in *Snca^NLS^* mice (left) with annotations for the dorsal and ventral horns and ventral white matter (right) from a section stained for αSyn. B) Merged micrographs from the dorsal horn (upper), ventral horn (middle), and ventral white matter (bottom) staining for neurons (left), astrocytes (middle left), microglia (middle right), and oligodendrocytes (right; different staining paradigm). Scale bars: 1,000 μm (A), 75μm (B).

**Supplemental Figure 5:** αSyn staining patterns throughout the spinal cord with neuron-specific markers for ChAT and Pax2. A) Coronal plane chosen to assess cervical (left) thoracic (middle) or lumbar (right) *Snca^NLS^* spinal cord in the intermediolateral (upper), lamina X (middle), and ventral horn (lower) co-stained for αSyn and ChAT. B) Coronal plane chosen to assess cervical (left) thoracic (middle) or lumbar (right) *Snca^NLS^* spinal cord in the dorsal horn (upper) and ventral horn (lower) co-stained for αSyn and Pax2. Scale bars: 75 μm.

**Supplemental Figure 6:** Mining scRNA-Seq datasets to explore αSyn expression patterns in the mouse spinal cord, brain, and colon. A) UMAP plot from the Harmonized Atlas of Mouse Spinal Cord with A’) *Snca* expression pattern and A”) neuron-specific sub-clustering. B) tSNE plot from the Aging Mouse Brain dataset from the Broad Institute’s Single Cell Portal with B’) *Snca* expression pattern and B”) neuron-specific sub-clustering from the study Dissecting Cell-Type Composition and Activity-Dependent Transcriptional State in Mammalian Brains by Massively Parallel Single-Nucleus RNA-Seq. C) tSNE plot from the Human and Mouse Enteric Nervous System at Single Cell Resolution from the Broad Institute’s Single Cell Portal with C’) *Snca* expression pattern and C”) neuron-specific sub-clustering. ABC (arachnoid barrier cells), ARP (astrocyte-restricted precursors), CC (colonocyte), CPC (choroid plexus epithelial cells), CSF-cN (cerebrospinal fluid contacting neurons), DC (dendritic cells), EPC (ependymocytes), Excit-(excitatory neurons), HbVC (hemoglobin-expressing vascular cells), HypEPC (hypependymal cells), Inhib-(inhibitory neurons), Inter-(interneurons), MN (motor neurons), MNC (monocytes), NendC (neuroendocrine cells), Neut (neutrophils), NRP (neuronal-restricted precursors), NSC (neural stem cells), OEG (olfactory ensheathing glia), OPC (oligodendrocyte precursor cells), PC (pericytes), PGC (preganglionic cells), Sensory (sensory neurons), Stem (stem cells), SV (secretomotor/vasodilator neurons), TA (transit-amplifying cells), TNC (tanycytes), VLMC (vascular and leptomeningeal cells).

## Graphical Abstract

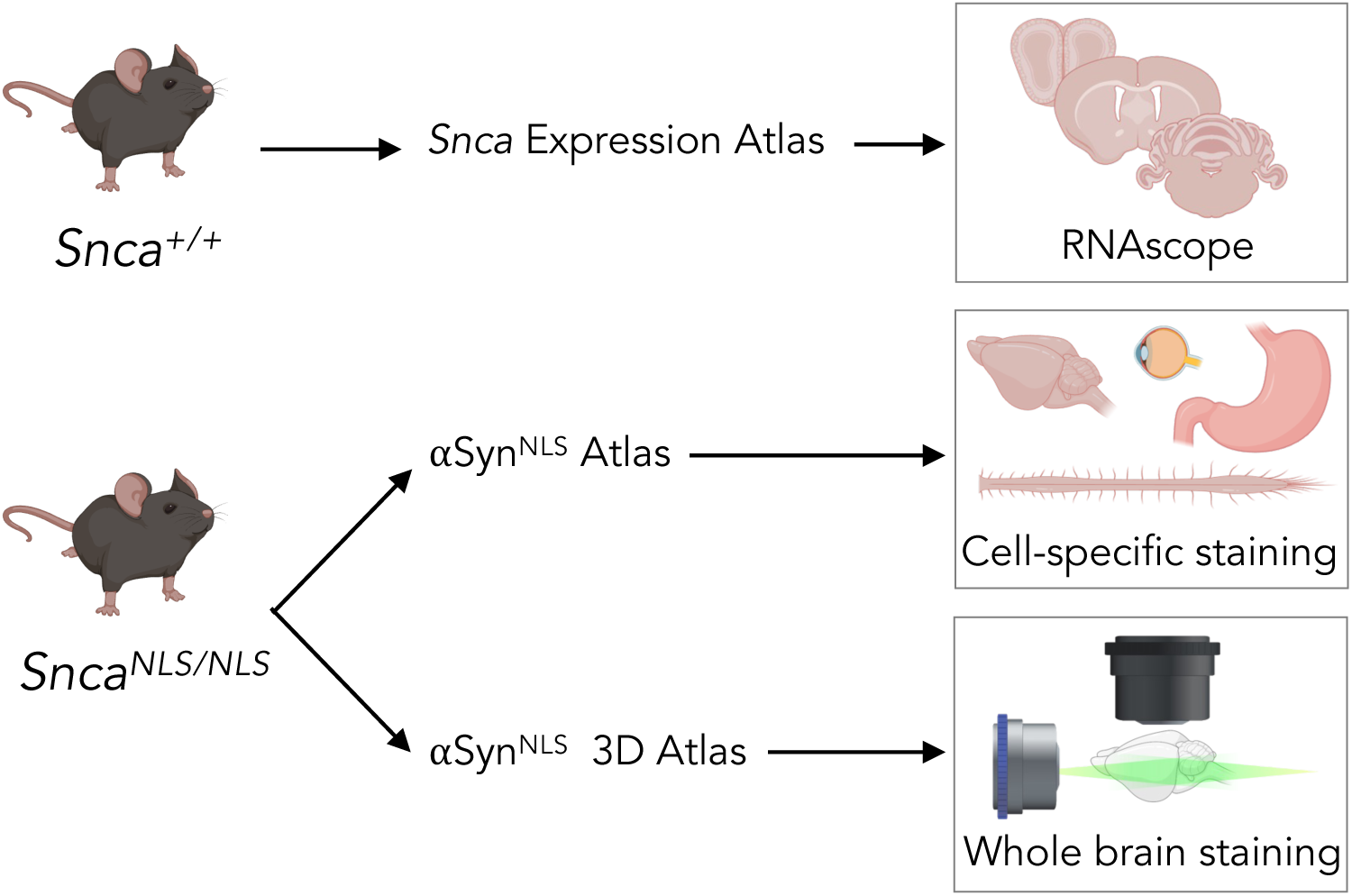

