## Supplemental Figures for "A topographical atlas of αSyn dosage and cell-type expression in the mouse brain and periphery"

A

### *Snca* Expression Atlas

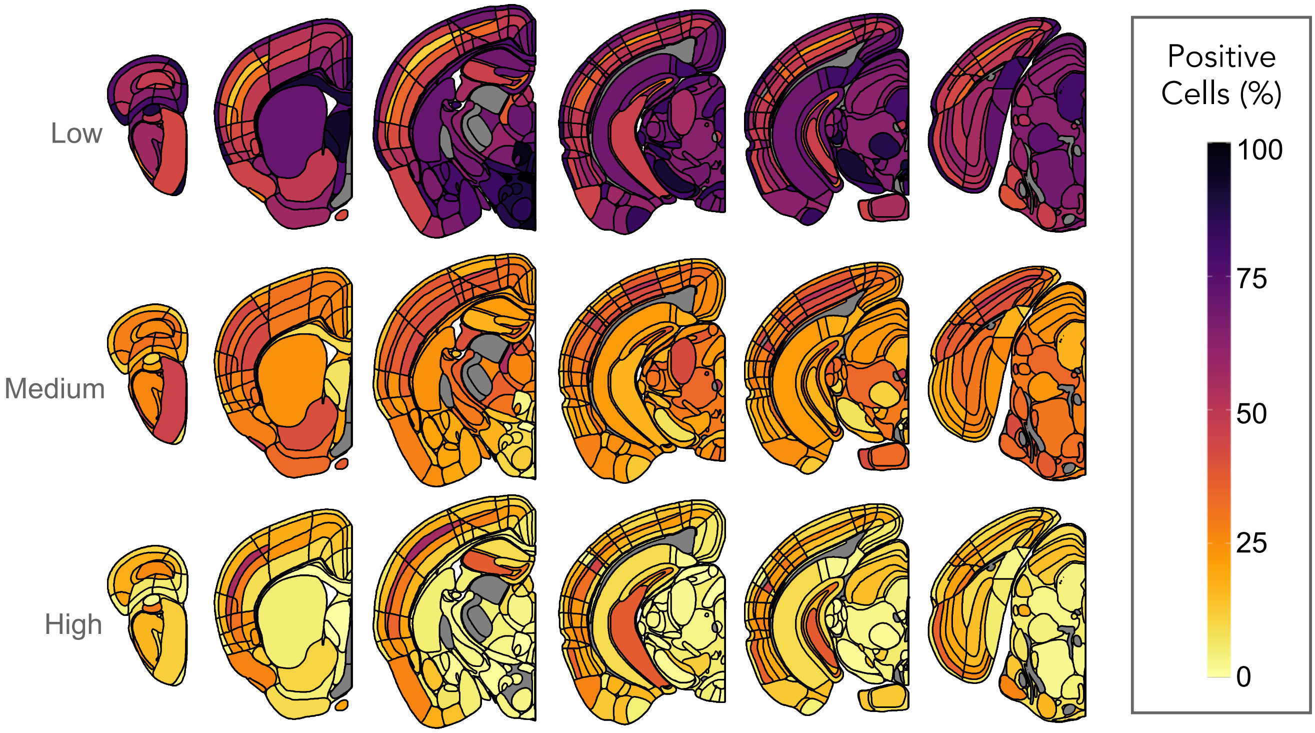


B

### αSyn$^{NLS}$ Atlas

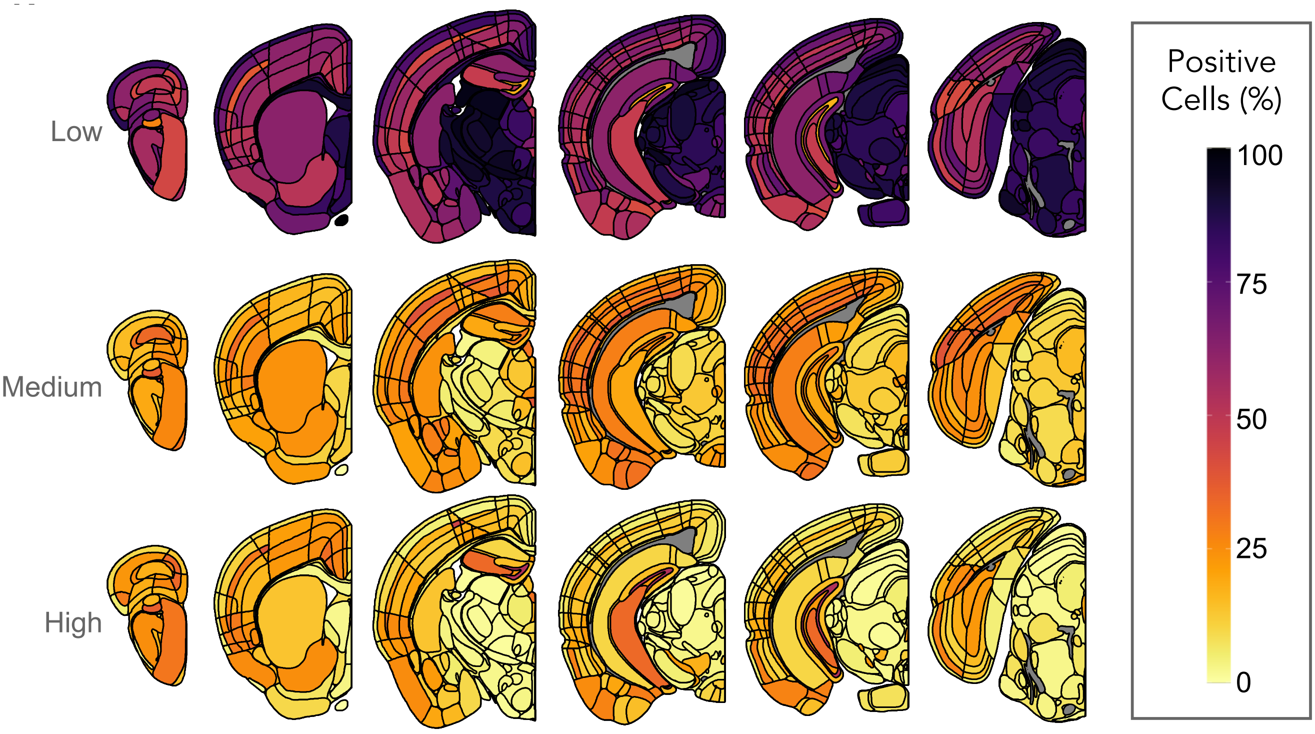


Supplemental Figure 1

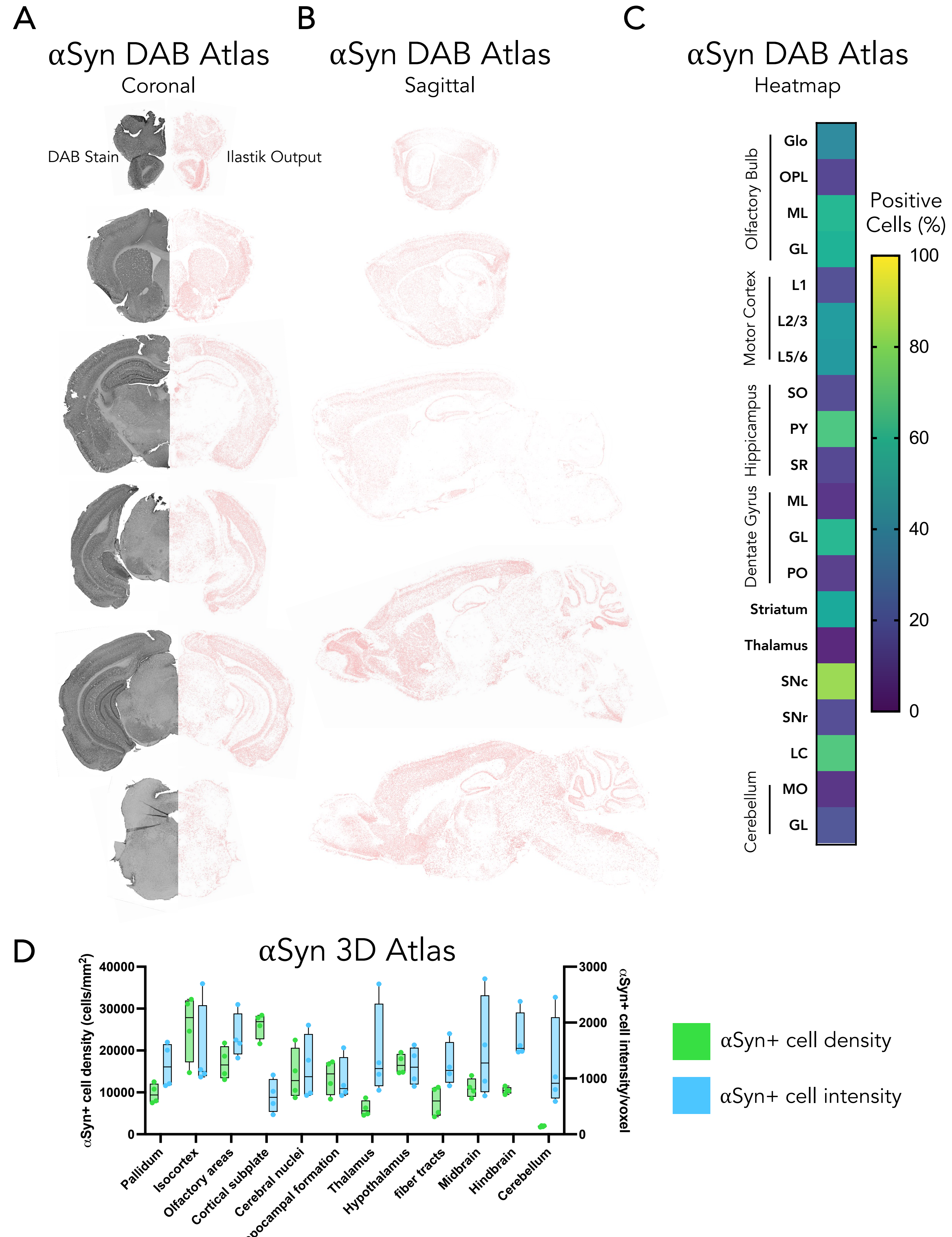


Supplemental Figure 2

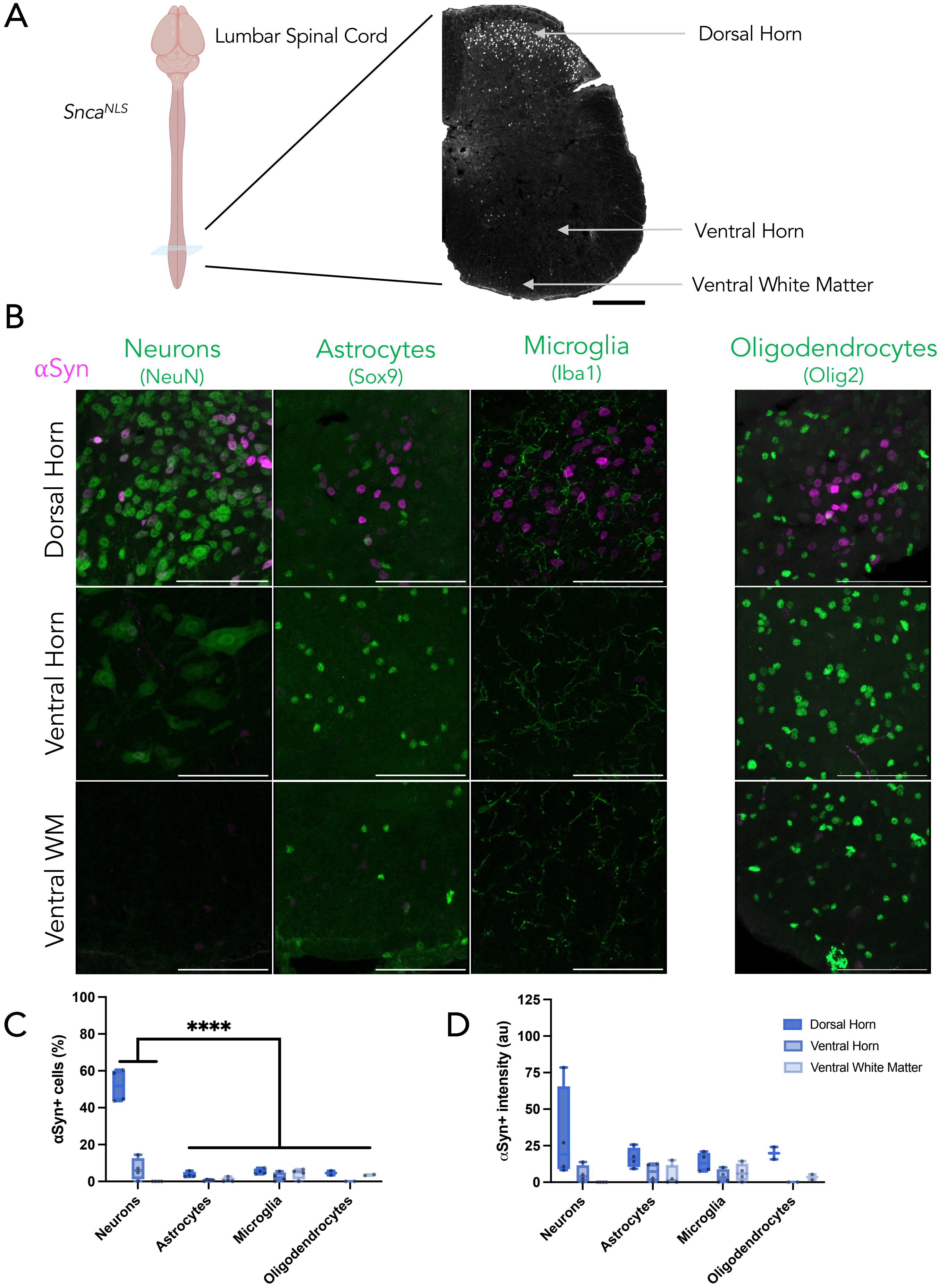


Supplemental Figure 3

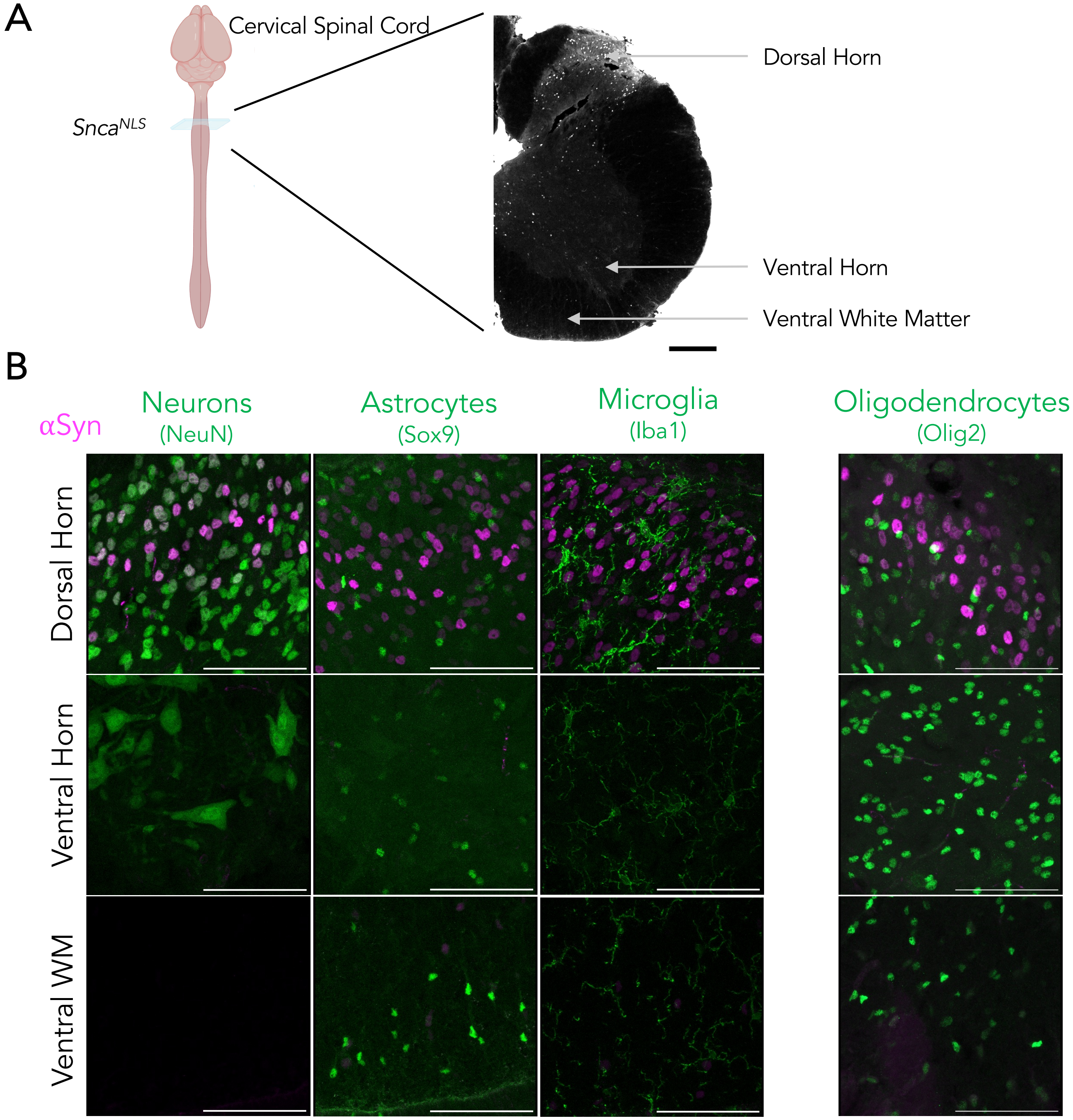
A
Cervical Spinal Cord
SncaNLS
Dorsal Horn
Ventral Horn
Ventral White Matter
B
αSyn
Neurons
(NeuN)
Astrocytes
(Sox9)
Microglia
(Iba1)
Oligodendrocytes
(Olig2)
Dorsal Horn
Ventral Horn
Ventral WM

A

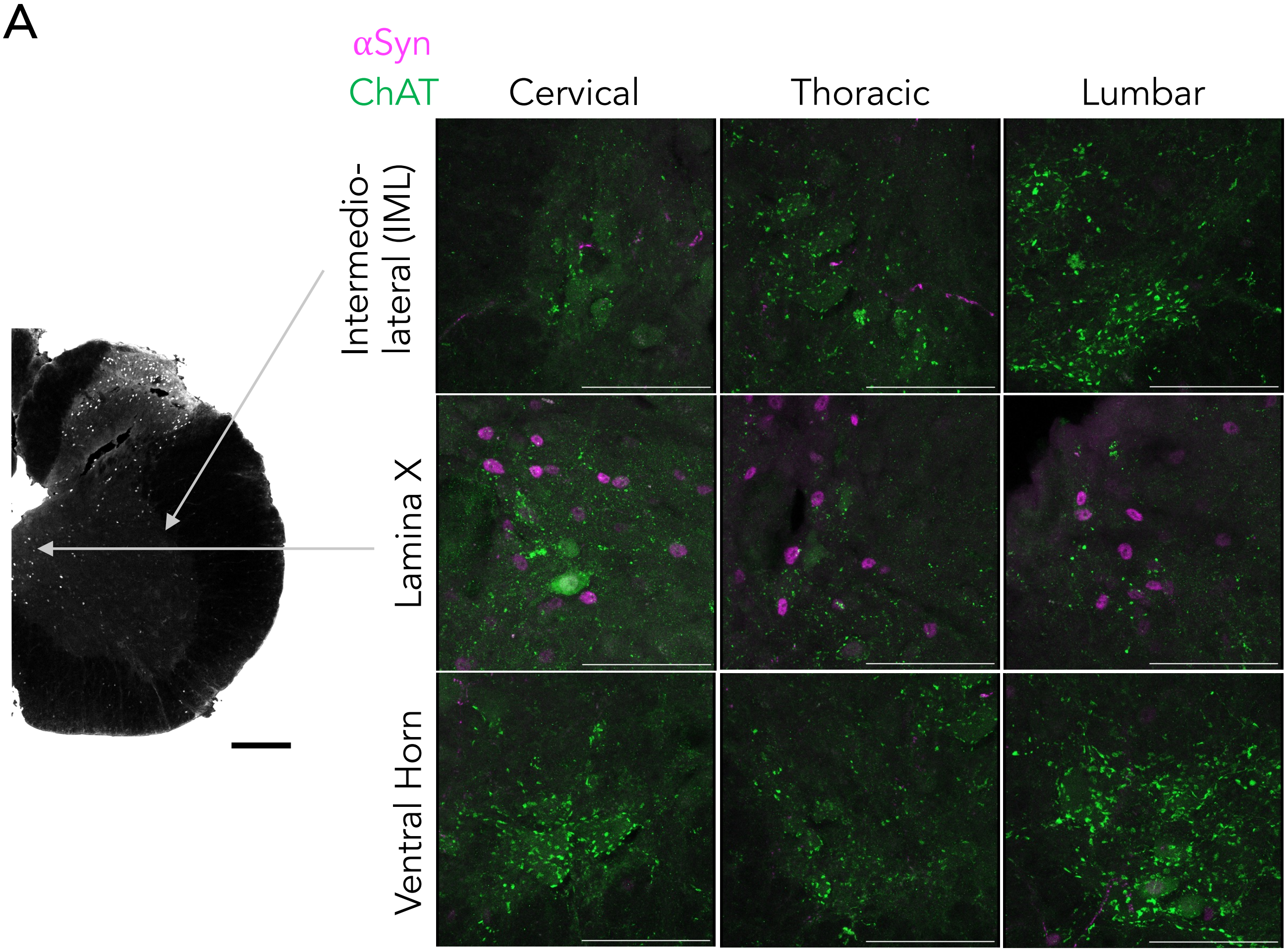


B

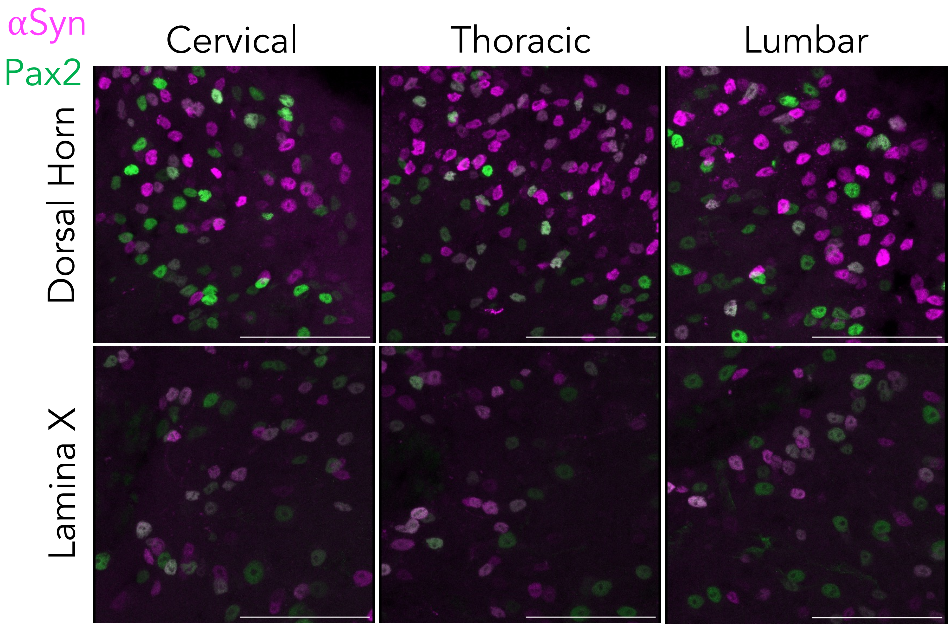


Supplemental Figure 5

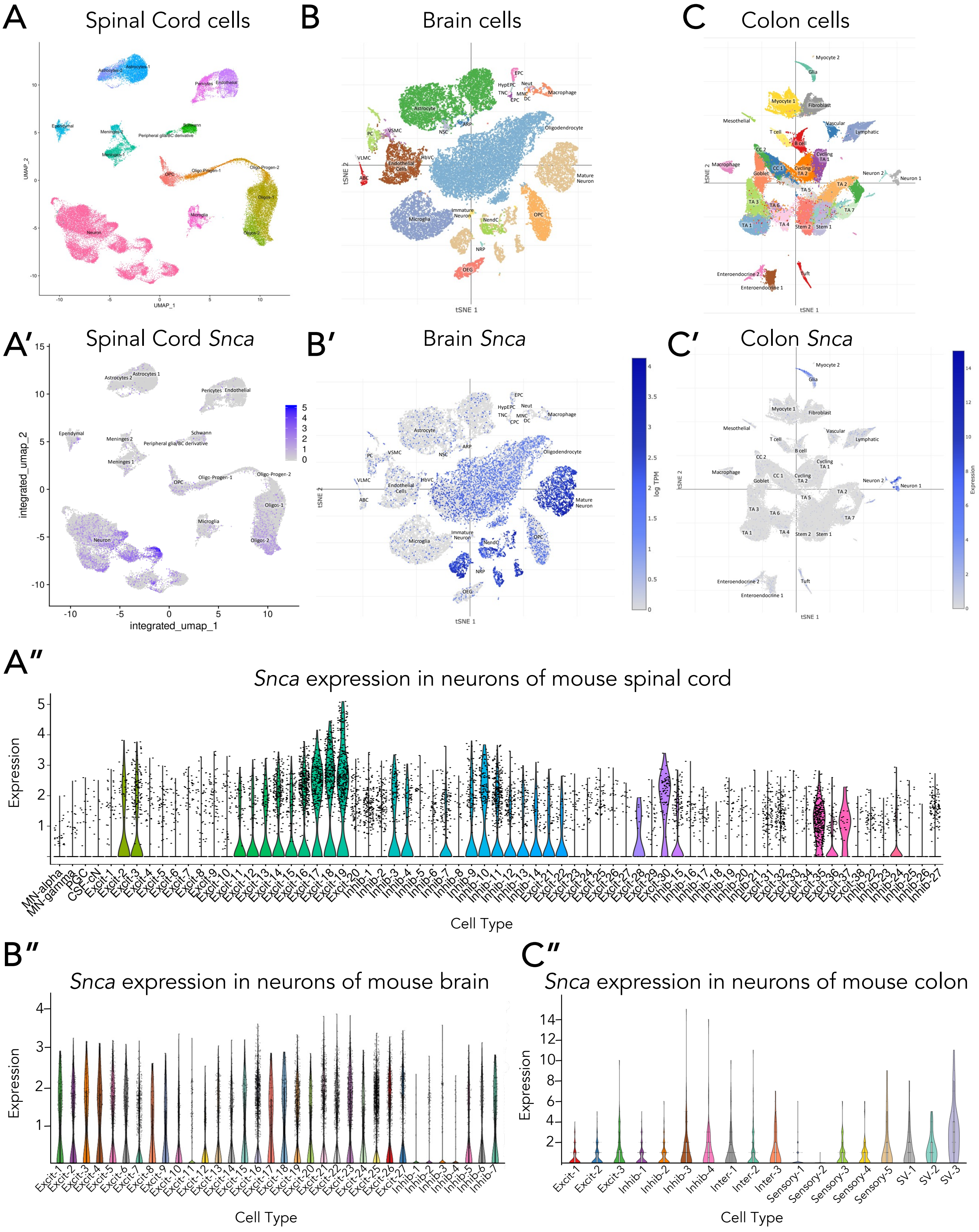


Supplemental Figure 6
